# Perivascular Excitation Tunnelling: a Novel and Preventable Cause of Cardiac Reperfusion Arrhythmias

**DOI:** 10.1101/2023.11.29.569031

**Authors:** Enaam Chleilat, Teo Puig Walz, Bo Han, T Alexander Quinn, Peter Kohl, Callum M Zgierski-Johnston

## Abstract

**Background:** Reperfusion after myocardial ischaemia can lead to deadly arrhythmias, in part due to heterogeneities in electrophysiology (EP) across affected tissue. There is a need to understand the spatiotemporal dynamics of ischaemia-reperfusion arrhythmias (IRA), so that reperfusion strategies to prevent them can be found.

**Methods:** Langendorff-perfused rabbit isolated hearts were loaded with a voltage-sensitive dye. Epifluorescence imaging was used to track action potential propagation across the cardiac surface. The heart was simultaneously perfused ‘globally’ (*via* the aorta) and ‘locally’ (*via* cannulation of a single coronary artery) with an oxygenated physiological saline solution. Local perfusion was subsequently switched to and from solutions that mimic aspects of ischaemia (acidosis, hypoxia, hyperkalaemia, or a simulated ischaemia solution combining all three) or to no-flow. Subsequently, different reperfusion strategies were tested to reduce IRA re-entries. The most successful strategy for preventing re-entry was tested in Langendorff-perfused isolated pig hearts to assess the clinical relevance of the observed mechanism and treatment strategy.

**Results:** Upon sudden reperfusion of the cannulated coronary artery in rabbit hearts we observed a preferential recovery of electrical excitability along the vessel’s main branch (‘perivascular excitation tunnelling’, PVET). This resulted in re-entry in roughly half of the hearts. Hyperkalaemia and hypoxia, but not acidosis, were sufficient to lead to conduction block, PVET, and re-entry, with both PVET and re-entry more frequently observed after hyperkalaemia than hypoxia.

PVET was also present in pigs and PVET-based re-entries were successfully prevented in rabbit and pig hearts by two-step reperfusion, first of the distal majority of the previously ischaemic region, and then of the remaining tissue from the proximal point. With this strategy, any PVET that developed in the distal tissue was blocked by the still inexcitable proximal tissue. Upon reperfusion of the proximal tissue, there was a reduced path length for PVET. As a consequence, the associated excitable gap was too short for re-entrant excitation.

**Conclusions:** We observed a novel arrhythmia mechanism upon coronary reperfusion (PVET), which suggests that preferential recovery of myocardial excitability along the reperfused vessel is an important mechanism underlying IRA formation. PVET-induced re-entry reliably occurred in both rabbit and pig hearts and could be prevented by two-step reperfusion.

**Graphical Abstract:** 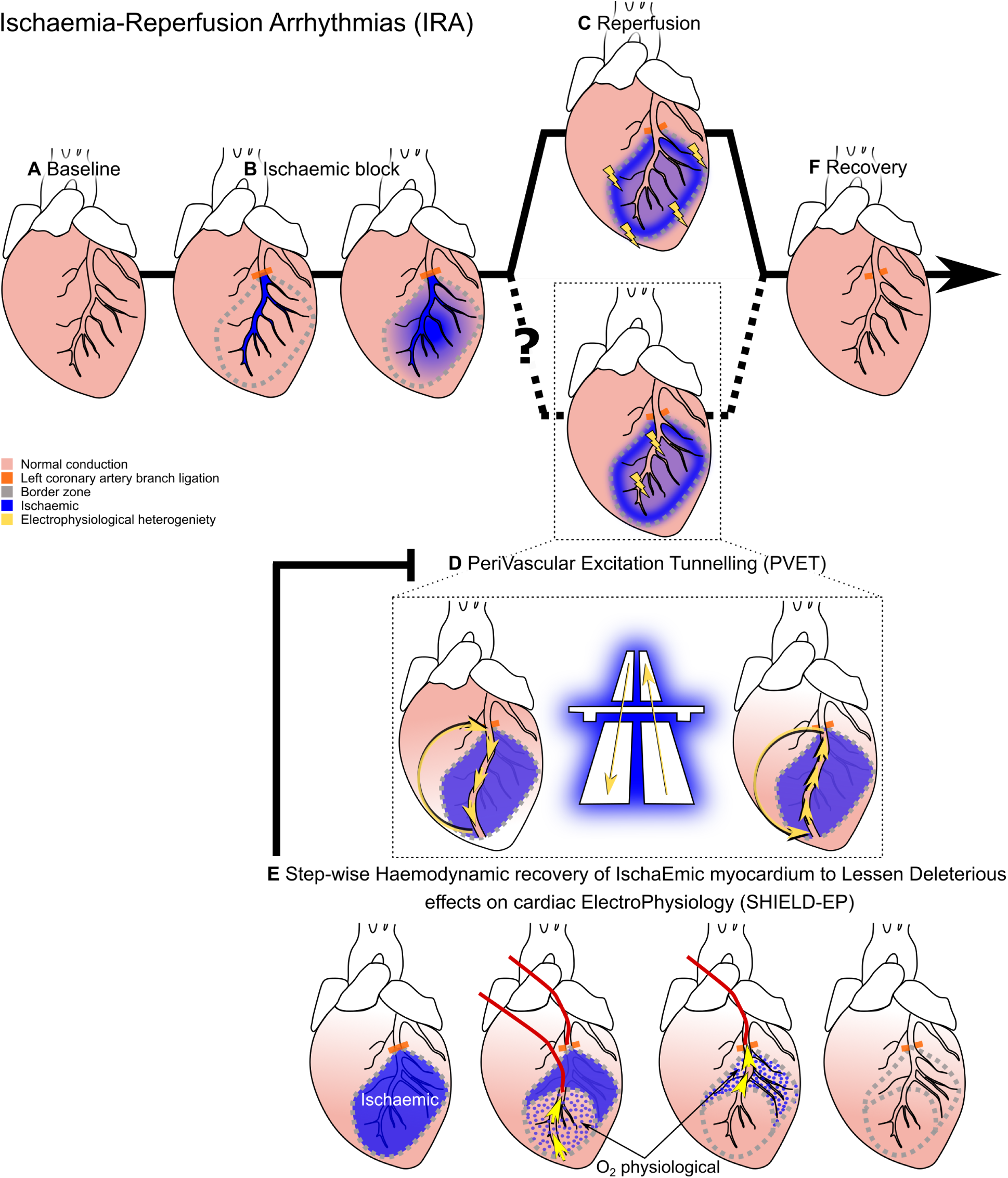

## Introduction

Cardiovascular diseases are the leading cause of incapacitation and death in the world.^1^ Morbidity and mortality from cardiovascular disease progression is often linked to cardiac arrhythmias. While mortality rates from arrhythmias have been declining, we are far from a mechanistic understanding of triggers and sustaining mechanisms underlying arrhythmogenesis.^2^ One of the currently unmet research problems is a comprehensive understanding of the spatiotemporal dynamics of electrophysiological (EP) changes in the context of myocardial ischaemia-reperfusion.

Myocardial ischaemia refers to inadequate coronary blood flow to (parts of) the myocardium, and its effects on cardiac function.^3^ The reduction in coronary blood flow can arise as a result of coronary atherosclerosis and/or -thrombosis, a possible treatment of which involves catheter-based removal of obstructions, and/or stenting of vessels (percutaneous coronary intervention; PCI). Typically, it takes 1-2 h from first symptoms to PCI.^4,5^ In the meantime, depletion of intracellular adenosine triphosphate (ATP), acidification, a decrease in extracellular and an increase in intracellular sodium concentration, and an increase in extracellular potassium concentration ([K^+^]_o_) and osmotic pressure in the affected tissue region contribute to altered EP properties, including detrimental effects on local excitability and conduction.^6^ Restoring blood flow (reperfusion) is critical to limit sustained myocardial damage. Paradoxically, though, reperfusion itself can give rise to fatal arrhythmias (ischaemia-reperfusion arrhythmias; IRA).^7^

While changes in the myocardial environment develop gradually after the onset of ischaemia, effects of reperfusion are sudden. This includes rapid restoration of near-normal physiological characteristics of the extracellular fluid. It has been suggested that this may increase EP heterogeneity in the ischaemic border zone due to washing of blood and extracellular fluid from central to peripheral ischaemic tissue,^8,9^ favouring arrhythmogenesis in that region.^10–19^

We hypothesise that, in addition to the above mechanism, IRA can occur as a result of the preferential restoration of the extracellular environment, and consequently myocardial excitability, in myocardium along the reperfused coronary artery. This may give rise to a preferential conduction pathway in the perivascular myocardium, which allows excitation tunnelling through the *centre* of the formerly ischaemic tissue. This ‘perivascular excitation tunnelling’ (PVET) acts as substrate for re-entrant arrhythmias during early reperfusion. Thus, we propose a new mechanism of IRA (the phenomenon), which we hypothesize is driven by PVET (the mechanism), and leads to re-entry (the consequence).

Assessing spatio-temporal dynamics of EP heterogeneity and arrhythmia formation *in vivo* is challenging, due to the low resolution of body surface electrocardiograms, the inability to spatiotemporally reliably contact-map large tissue areas with catheter-based systems, and the fact that data obtained with either approach originate from extracellular signals. *Ex vivo* epifluorescence imaging with voltage sensitive dyes (‘optical mapping’; OM) in Langendorff-perfused hearts allows for high spatio-temporal resolution recordings of transmembrane voltage in a well-controlled setting.^20^ This control includes not only the option of simulating ischaemia as global or local ‘no flow’, but also the possibility of breaking this down into individual ‘ischaemic components’, such as hyperkalaemia (elevated [K^+^]_o_; HiK^+^), hypoxia (reduced oxygen partial pressure, potentially combined with metabolic inhibition, for example by 2-deoxy-D-glucose and sodium cyanide; LoO_2_), or decreased pH (acidosis; HiH^+^). Individually modifying these factors may allow identification of key mechanisms underlying PVET, and the development of innovative reperfusion strategies that mitigate re-entry.

The aim of the current study is to elucidate in isolated heart experiments: (i) the presence of PVET; (ii) its potential contribution to IRA; (iii) potential contributors (HiK^+^, LoO_2_, HiH^+^); (iv) whether ‘smart’ reperfusion strategies may prevent PVET-associated re-entries; and (v) if results are translationally relevant in a pig model. Our main hypotheses are: (i) reperfusion of locally ischaemic tissue is associated with pronounced EP heterogeneity not only at the border between ischaemic and control tissue, but also along the main trunk of the reperfused coronary artery in the central ischaemic zone; (ii) this can give rise to the formation of preferential conduction pathways in perivascular myocardium, which may support re-entrant excitation by PVET; (iii) PVET is caused predominantly by the swift normalisation of [K^+^]_o_; and (iv) PVET can be avoided by reperfusing ischaemic tissue in a controlled two-step manner in which recovery of excitability is split into two stages (distal-first, proximal-last) with the inexcitable proximal tissue preventing re-entry.

## Methods

### Animals

Experiments were performed in both rabbit and pig hearts. New Zealand White rabbits were used for this study: 4-month-old (*N*=66 (animals), *n*=105 (reperfusions)) and 2-month-old (*N*=31) rabbits (strain code 052, Charles River Laboratories, Romans, France). Male or female rabbits were used indiscriminately based on availability (larger rabbits: 42 ♀ and 23 ♂; smaller rabbits: 29 ♂ and 2 ♀) with body weight at 3155 ± 47 g and a heart tissue weight of 9.58 ± 0.23 g for the larger cohort *vs.* 1473 ± 42 and 5.98 ± 0.24 g for the smaller animals (Fig. 5). Munich hybrid minipig^21^ (6-months; courtesy of Prof Angelika Schnieke, TUM, Germany) were used at ∼50 kg body weight for the translational validation (*N*=16; 15 ♀ and 1 ♂ pigs).

All investigations reported in this manuscript conformed to German animal welfare laws (TierSchG and TierSchVersV), compatible with the guidelines stated in Directive 2010/63/EU of the European Parliament on the Protection of Animals used for Scientific Purposes, and were performed with ethical approval by the local Institutional Animal Care and Use Committee (Regierungspräsidium Freiburg, X-16/10R and X-21/06R for rabbits and X-21/03B for pigs). Animal housing and handling adhered to good animal practice, as defined by the Federation of European Laboratory Animal Science Association (FELASA).

### Heart Preparation

Rabbits were anaesthetised using a mix of ketamine (0.5 mL/kg) and xylazine (0.2 mL/kg) injected intramuscularly (i.m.) in the hindlimb, then heparinized (1,000 units/rabbit) and euthanized by sodium thiopental overdose (40 mg/kg body weight) via ear vein injection. After thoracotomy, the heart was swiftly excised and perfused (<2 min from excision) with oxygenated HEPES-buffered Tyrode’s solution (HT, see Table 1) at a rate of 17.5 mL/min. An incision into the proximal pulmonary artery allowed coronary effluent to exit the right ventricle. Remaining non-cardiac tissue (lungs, thymus, pericardium, vessels) was removed.

**Table 1:** Composition of HEPES-buffered Tyrode’s (HT) solution, simulated ischaemia (SI) solution, and solutions mimicking (HiK^+^), hypoxia (LoO_2_), and acidosis (HiH^+^) in larger rabbit experiments. NaOH was used to titrate pH at 37°C.

| Reagent | Concentration (mmol/L) |  |  |  |  |
| --- | --- | --- | --- | --- | --- |
|  | HT | SI | HiK<sup>+</sup> | LoO<sub>2</sub> | HiH<sup>+</sup> |
| NaCl | 140 | 140 | 140 | 140 | 140 |
| KCl | 6 | 15 | 15 | 6 | 6 |
| HEPES | 10 | 10 | 10 | 10 | 10 |
| Glucose | 10 | - | 10 | - | 10 |
| 2DG | - | 20 | - | 20 | - |
| CaCl<sub>2</sub> | 1.8 | 1.8 | 1.8 | 1.8 | 1.8 |
| MgCl<sub>2</sub> | 1.0 | 1.0 | 1.0 | 1.0 | 1.0 |
| NaCN | - | 1.0 | - | 1.0 | - |
| pH | 7.4 | 6.5 | 7.4 | 7.4 | 6.5 |
| Gas for bubbling | O<sub>2</sub> | N<sub>2</sub> | O<sub>2</sub> | N<sub>2</sub> | O<sub>2</sub> |
| Osmolality (mOsm/kg) | 300 ± 5 | 320 ± 5 | 308 ± 0.4 | 313 ± 0.7 | 293 ± 0.5 |

2DG: 2-deoxy-D-glucose; NaCN: sodium cyanide. Composition of SI after Khokhlova et al.<sup>23</sup>

For the pigs, sedative premedication was used at 0.5 mg/kg body weight midazolam and 20 mg/kg body weight ketamine i.m. (mixing syringe). This was followed by i.v. (ear vein) administration of 2-4 mg/kg propofol. Euthanasia was carried out by lethal dose of propofol (20-60 mg/kg) or a lethal dose of KCl (20-40 mL 7.45%) in deeper anesthesia (*i.e.* after propofol). It took an average of 3.5 minutes from the first skin incision to heart removal. The particularly critical time from heart removal to coronary perfusion was <1.5 minutes.

### Local Perfusion and Panoramic Optical Mapping of Rabbit Hearts

Panoramic, transepicardial, 3-view OM recordings were obtained for high-resolution spatio-temporal visualisation of transmembrane potentials in isolated Langendorff-perfused rabbit hearts (Fig. 1). Hearts were connected to a modified Langendorff perfusion system (Hugo Sachs Elektronik, March, Germany; Fig. 1), monitoring aortic pressure (maintained at 80 mmHg), flow rate, temperature, and pseudo-ECG (monopolar Ag/AgCl pellet electrodes on the cardiac surface). Hearts were loaded with voltage-sensitive dye (25 µL bolus of 8.6 mM di-4-ANBDQPQ, Cytocybernetics, Buffalo, NY, USA) and electromechanically uncoupled ((±)blebbistatin ([Abcam, Amsterdam, Netherlands] or [Biozol, Eching, Germany] or [TRC, Toronto, Canada] or [Tocris, Bristol, UK], at final concentration 10 µM).

**Figure 1:**
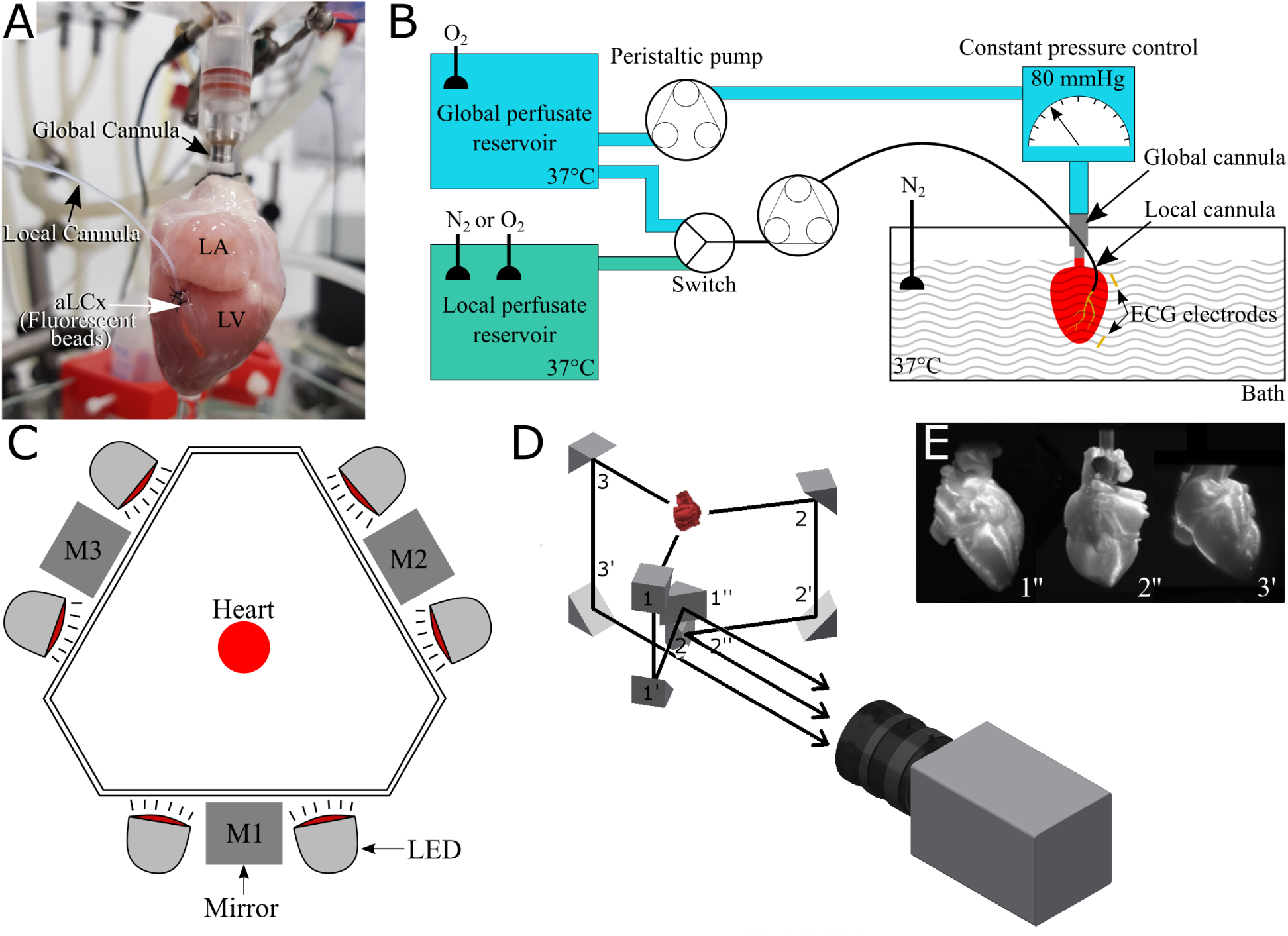
Experimental setup. (A) Isolated rabbit heart, perfused in Langendorff mode via the global cannula, and simultaneously perfused locally via a cannula (local cannula) in the anterior branch of the left circumflex coronary artery (aLCx). (B) Schematic illustration of the setup. Global Langendorff perfusion (at 80 mmHg) and local aLCx perfusion (at ∼8 mL/min) are individually controlled; local perfusion can be switched to no-flow, or to perfusates mimicking distinct aspects of myocardial ischaemia (for detail, see Table 1). (C) Birdseye schematic view of bath, mirrors (M#) and LED positions. (D) Schematic illustration of the equidistant path lengths for panoramic projection of three views of the heart onto a single sCMOS camera. (E) Optical projections of a 3D-printed mock rabbit heart obtained with this setup. Note: in (A), red fluorescent beads were injected at the end of a study into the aLCx to visualise the topology of locally perfused vasculature.

The three optical mapping views were projected onto a scientific Complementary Metal–Oxide– Semiconductor (sCMOS) camera (Andor Zyla 5.5, Oxford Instruments, UK) recording at a frame rate of 100-200 Hz. Eight band-pass filtered (ZET642/20X, Chroma Technology, Bellows Falls, VT, USA) red LED (CBT-90R, Luminus, Sunnyvale, CA, USA) were used to excite the dye, with fluorescence emission collected *via* a high-speed lens (DO-5095, Navitar, Rochester, NY, USA) and a custom filter (ET585/50M+800/200M, Chroma). The fluorescence signal was analysed using MATLAB (Mathworks, Natick, MA, USA; routines available upon request from the authors). The background of the images was masked and the remaining signal inverted (to mimic usual action potential recordings) and normalised on a pixel-by-pixel basis using following formula:

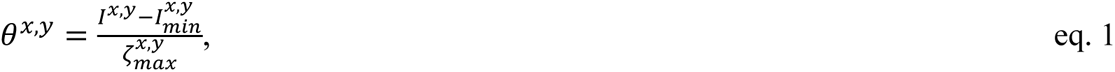

where the normalised intensity for each pixel, 𝜃^𝑥,𝑦^, was computed by subtracting from each pixel value, 𝐼^𝑥,𝑦^, the baseline of the measured intensity for that pixel, 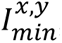, and normalising by the maximum intensity measured on that pixel in a reference recording with control conduction through the myocardium, 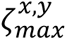. This reference-normalisation avoids the enhancement of low-level signals in conduction-blocked regions which would be artificially increased by normalising with the local maxima in recording without cells undergoing excitation. Subsequently, a spatial median filter with a kernel size of 3 × 3 was applied on each image. Spatially averaged signals were computed for three regions of interest (∼20 × 20 pixels), representing atrial, globally-perfused ventricular, and locally perfused ventricular tissue. To compute activation-time maps, ensemble averaging was performed over a 5 second recording after spatial filtering was applied, and activation-time on a per-pixel basis was calculated as the maximum of the interpolated signal time derivative (*e.g.* Suppl. Fig. 2B).

In human, myocardial ischaemia is most frequently a result of blockage (thrombosis) caused by rupture of an atherosclerotic plaque of the left anterior descending coronary artery. In rabbit, the functional equivalent is the anterior branch of the left circumflex coronary artery (aLCx; though generally, the left anterior descending coronary artery is a relatively minor vessel).^22^ The aLCx was cannulated to allow controlled local perfusion of a region of the left ventricle. To do so, two sutures (0.07 mm Silkam® silk, braided, coated, non-absorbable sutures; Ref C0764043; 6/0 B. Braun Deutschland GmbH & Co.KG, Melsungen, Germany) were passed under the aLCx (10 mm ½ circle round body atraumatic needle) at with ∼2 mm between sutures. The aLCx surface was then cut open between sutures, and a custom-made cannula (inner diameter 0.3 mm; outer diameter 0.6 mm, Ref Z609692, Sigma-Aldrich Chemie GmbH, Taufkirchen, Germany) was inserted into the aLCx. The proximal suture was used to ligate the truncated end of the aLCx, the distal suture sealed the vessel around the cannula, and both sutures secured the local perfusion line in place. The perfusate used for local perfusion was initially identical to the global HT perfusate, with flow adjusted to match pre-cannulation coronary flow (using the global flow to predict the local flow rate; ∼8 mL/min *via* the aLCx). Local perfusion was then switched either to and from no-flow, or to solutions that simulate certain aspects of myocardial ischaemia (HiK^+^, LoO_2_, HiH^+^, or a combination of all three; simulated ischaemia solution, SI; see Table 1).

Partial pressure of oxygen (*p*O_2_) in the perfusates was monitored during experiments. Physiological perfusates were O_2_-bubbled and had a *p*O_2_ of 674 ± 19 mmHg (*n*=31); LoO_2_ and SI perfusates were N_2_ bubbled (see Table 1) and had a reduced *p*O_2_ of 32 ± 13 mmHg (p<0.0001, *n*=22) and 44.82 ± 8, n=22 respectively. Non-bubbled solution had a *p*O_2_ of 175 ± 23 mmHg, which exceeds *p*O_2_ levels in arterial blood (∼50 mmHg). The bath surrounding the heart was therefore also N_2_-bubbled. As oxygenated perfusate from the global solution enters the bath, which furthermore has a large interface with room air, bath *p*O_2_ was 126 ± 11 mmHg (*n*=24).

For each experiment, we first recorded normal baseline electrical excitation. Irregularities in excitation patterns due to technical (*e.g*., inadequate perfusion *via* the local cannula or insufficiently homogeneous dye loading) or biological problems (*e.g*., damage to the preparation or cardioversion inability out of baseline ventricular fibrillation / tachycardia) formed exclusion criteria. Once baseline activity was established (example in Fig. 2A), we switched local perfusion to no flow or one of the solutions mimicking ischaemic factors and recorded OM once per min. Local perfusion was switched back to physiological solution 2 min after block was detected. If conduction block did not occur, perfusion was switched back to physiological solution after ∼1 hour. Upon reperfusion, 5 min long OM recordings were obtained to catch any PVET or other arrhythmias.

**Figure 2:**
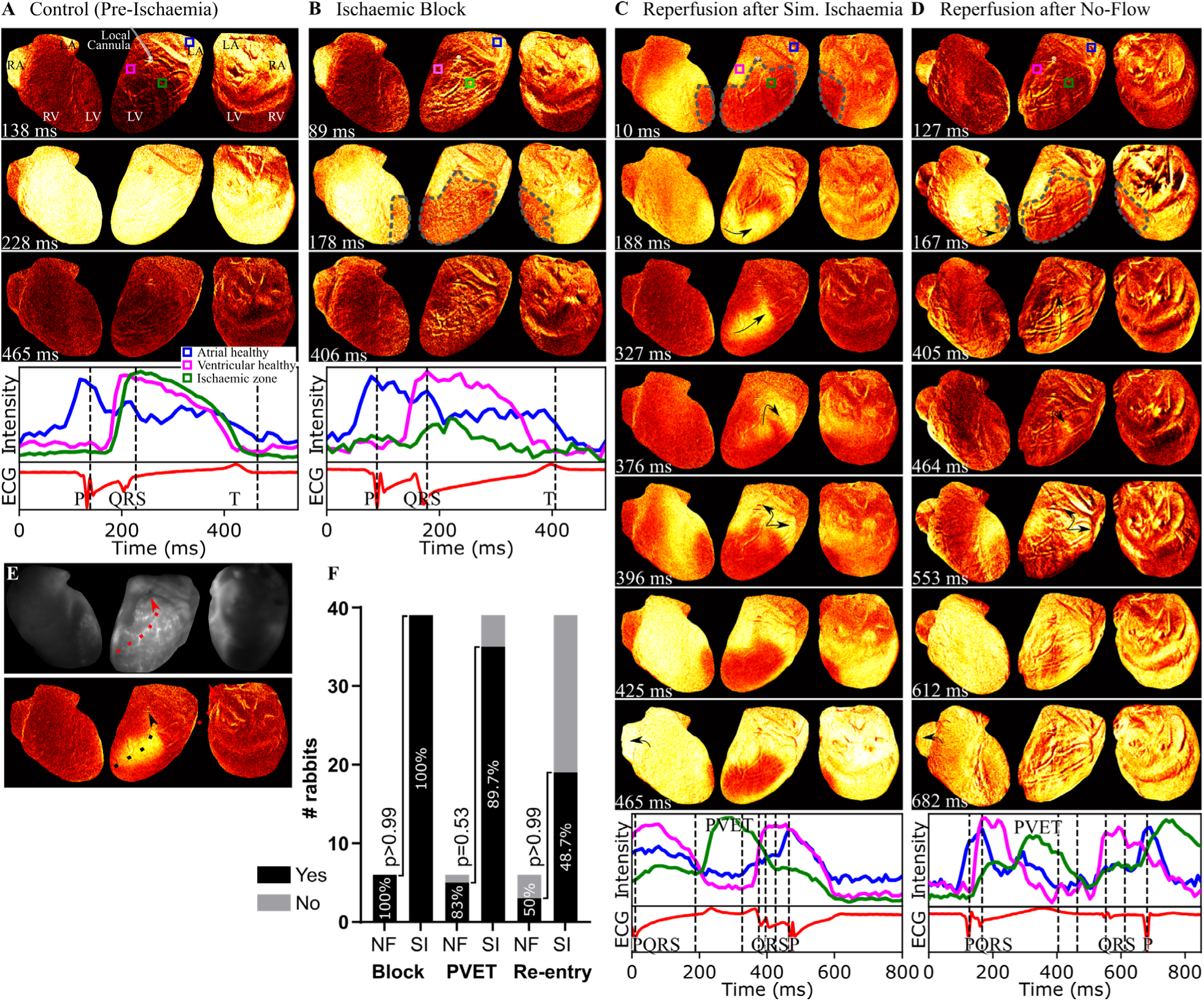
Example of PVET in *ex vivo* rabbit heart. In OM frames, intensity of transmembrane potential swings from most negative to most positive values are intensity-coded. In the time-plots, pseudo-ECG recorded from the cardiac surface (red trace) and action potentials optically recorded from from three regions of interest (blue = left atrium, pink = right ventricle away from the locally perfused tissue, green = locally perfused left ventricle) are shown. (A) Physiological activation of the heart after local cannulation, during perfusion of the entire heart using oxygenated physiological saline. (B) Local block of conduction (dashed outline), (C) PVET upon reperfusion after 2 min of local perfusion with simulated ischaemia (SI). Repeat-interventions were interspersed by 15 min normal perfusion. (D) PVET upon reperfusion, following 54 min of regional no-flow (NF; black arrows indicate the direction of activation-waves). (E) After termination of the experiments, fluorescent beads were washed in through the local perfusion line, showing the sub-epicardial vascular bed morphology of the target area (red arrow indicates the path of ascending PVET observed in C; note that the position of the heart relative to the camera changed slightly between runs). (F) Summary of incidences of block, PVET and PVET-based re-entries for no flow experiments (n=6) and for simulated ischaemia (n=39).

Inter-individual variability in coronary architecture meant that the locally perfused area differed between rabbits. The three-view mirror-based panoramic system with equal path lengths between each part of cardiac surface and the cameras ensured that the perfused tissue was always in view (Fig. 1).

### Large Animal Optical Mapping

Isolated pig hearts were initially Langendorff perfused by gravity, using HT (to wash out blood) then cardioplegic solution for transport from the operating theatre to our laboratories. In the optical mapping suite, a constant pressure Langendorff system was used to perfuse the heart with HT + 10 mM 2,3-butanedione monoxime (BDM; Sigma-Aldrich Chemie GmbH, Taufkirchen, Germany) solution. The aLCx, also known as left marginal artery in larger mammals, was cannulated and independently perfused (Fig. 7a’-a’’). Initially, all tissue was perfused with HT. Then, regional myocardial ischaemia was simulated by directing N_2_-bubbled HiK^+^ to the local perfusion line. The best reperfusion approach uncovered in rabbits was tested in pigs.

### Coronary Morphology Assessment

Information on coronary flow dynamics was collected in rabbits by perfusing the cannulated coronary artery with a ∼7 µL bolus of 0.25 mM Dextran, Alexa Fluor 680 (molecular weight 3 kDa, Stoke’s radius 0.95 nm, anionic, CAT.NO D34681, Thermo Fisher Scientific, Darmstadt, Germany). The spread of fluorescence within the tissue was monitored using a MVX10 macro zoom microscope (Olympus Life Science, Tokyo, Japan) and the same sCMOS camera and illumination/collection optics as for OM. Following OM experiments in pig hearts, the fluorescent dextran was locally perfused to visualise aLCx topology directly from the cMOS.

Following each OM experiment in rabbit hearts, red 6 µm diameter FluoroMax fluorescent polymer microspheres (CAT.NO 36-2, Thermoscientific, CA, USA) were injected through the local cannula to visualise the perfusion bed.

### Reagents

### Statistics

Fisher’s exact test was used for assessing significance (GraphPad Prism 7.05). Data was normalized to % of total included experiments to compare across conditions / reperfusion strategies. Unpaired t tests were used for comparison between small and large hearts.

## Results

### Local Ischaemia-Reperfusion *Ex vivo*: No-Flow and Simulated Ischaemia

In rabbit experiments (Fig. 1), local no-flow (NF) ischaemia and perfusion with SI solution both led to regional conduction block (Fig. 2B) of the previously normal cardiac conduction (Fig. 2A). The only observed difference between local no-flow ischaemia and local simulated ischaemia was in the time-frame to reach conduction block, with the no-flow approach requiring ∼45 min and simulated ischaemia ∼5 min.

Upon reperfusion from NF ischaemia, we observed formation of an excitable path in the myocardium along the cannulated aLCx (PVET; Fig. 2D; 5 of 6 hearts); this also occurred after local perfusion with SI (Fig. 2C; 35 of 39 hearts). This resulted in a period of heterogeneous excitability with slowed conduction entering and traversing the predominantly non-excitable ischaemic region. In Fig. 2C-D the excitable path follows the apico-basal direction, giving rise to a premature ventricular and retrograde (ventricular-atrial) conduction. Reperfusion in both cases led to complete recovery of conduction within 10 - 15 min of reperfusion. During no-flow-reperfusion, 83.3% of hearts produced PVET and 50.0% re-entry (Fig. 2F, *N*=6). During SI-reperfusion, 89.7% exhibited PVET and 48.7% manifested re-entries (Fig. 2F, *N*=39; 6 hearts were excluded due to technical issues and 1 heart excluded as arrhythmia occurred before reperfusion). There was no statistical significant difference between NF and SI (block p>0.9999; PVET p=0.5287; re-entry p>0.9999 by Fisher’s exact test) so for the remainder of the rabbit experiments, SI was used to more rapidly model ischaemia.

Following OM, red fluorescent beads were washed-in through the local cannula to visualise the perfusion bed (Fig. 2E). Comparing aLCx morphology with excitable tunnel topology, we found that preferential early restoration of excitability occurred along the main trunk of the reperfused coronary artery. In line with these findings, experiments on coronary flow dynamics illustrate that the myocardium closest to the main coronary artery trunk is perfused earlier than more distant tissue (Fig. 3).

**Figure 3:**
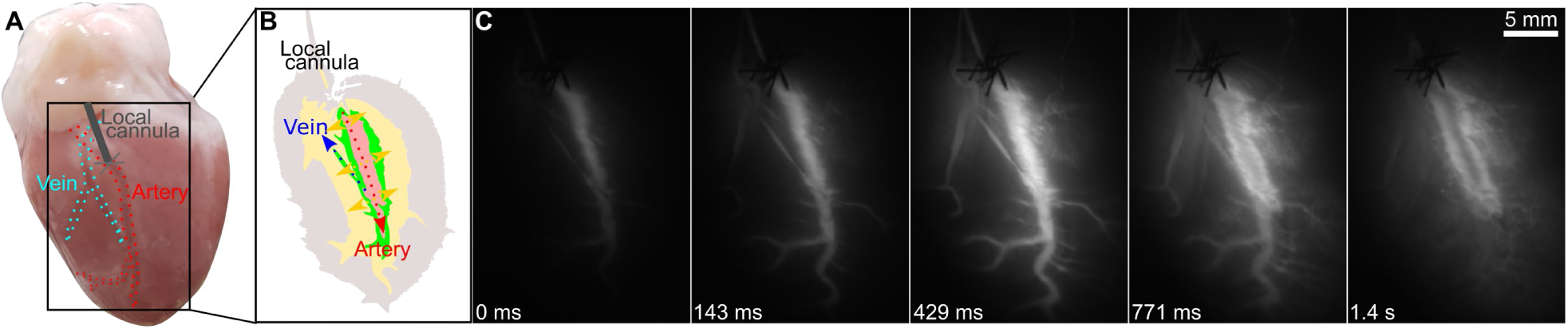
Visualisation of spatial dynamics of myocardial tissue perfusion upon acute changes in coronary artery content. Overview of rabbit heart (A), highlighting a branch of the aLCx and a nearby vein. (B) Threshold based schematic of lateral spread of fluorescent in (C). Bolus application of fluorescent dextran (Stoke’s radius 0.95 nm) into the aLCx, starting at 0 ms, leads to a rapid increase in fluorescence in the main trunk of the coronary artery (143 ms). This is followed by increased fluorescence in more distant sections of the vascular tree (including the vein seen in A-B: blue arrows). Interestingly, there is pronounced lateral spread of fluorescence from the main coronary artery into the perivascular myocardium (429 ms - 1.4 s; yellow arrows). As dextrans are small, they enter the extracellular space where they diffuse. This observation illustrates a preferential wash-in of fluorescence into the perivascular space.

### Experimental Drivers of PVET

Hearts were perfused with different solutions mimicking the different aspects of ischaemia (hyperkalaemia, hypoxia, acidosis). Block (Fig. 4B, F), PVET (Fig. 4C, G), and re-entry occurred when perfusing with either HiK^+^ (Fig. 4C) or LoO_2_ (Fig. 4G) solution. Altogether, during HiK^+^ (Fig. 4I, *N*=10), conduction block occurred in 80% of hearts, which resulted in PVET in all hearts and 60% of which showed re-entry. During LoO_2_, block occurred in all hearts, which resulted in PVET in 40% and 30% of which showed re-entry (Fig. 4I, *N*=10). Acidosis caused a minor slowing in activation but no block and therefore no PVET or re-entry (Fig. 4I, Suppl. Fig. 2, *N*=12). There was no significant difference between SI and HiK^+^ except for the likelihood of conduction block (*p*=0.0383). A high prevalence of PVET and re-entry exists in both SI and HiK^+^. When comparing SI and LoO_2_, there were fewer PVET events with LoO_2_ (*p*=0.0022), but the incidence of block or re-entry were not different. SI and acidosis significantly differed in both the incidence of block and PVET (*p*<0.0001), as well as re-entry (*p*= 0.0017). Incidence of all EP changes were not significantly different between HiK^+^ and LoO_2_. HiK^+^ compared to HiH^+^ was different in the incidence of block (*p*=0.0001), PVET (*p*=0.0001), and re-entry (*p*=0.0028). LoO_2_ was different from HiH^+^ in the incidence of block (*p*<0.0001) and PVET (*p*=0.0287), but not re-entry (*p*= 0.0779).

**Figure 4:**
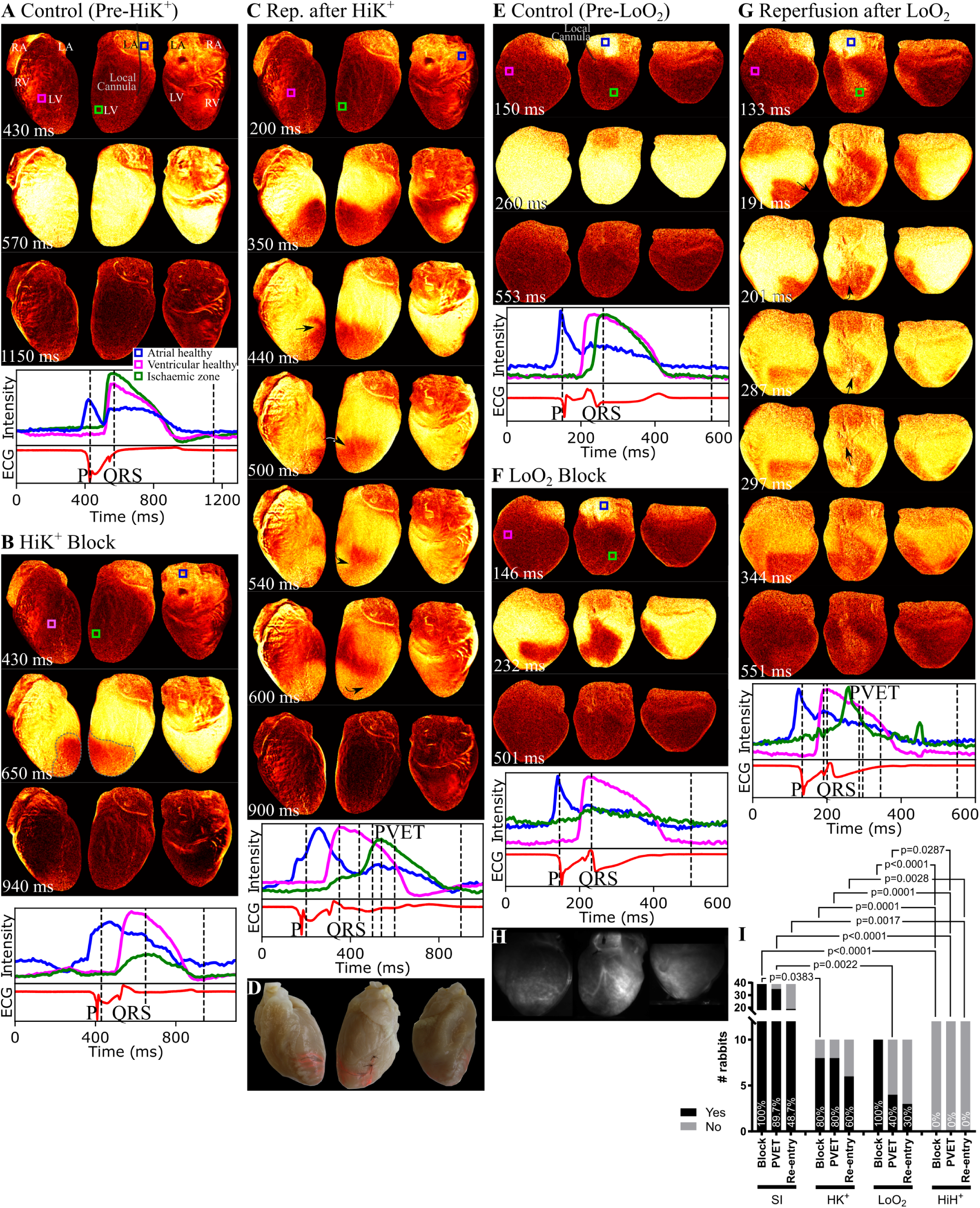
**Hyperkalaemia and hypoxia as main aspects driving PVET**. In a cohort of heart, hyperkalaemia and hypoxia (not acidosis) were the significant players driving arrhythmias as suggested by modelling. (A) Optical mapping during sinus rhythm pre-hyperkalaemia. (B) Optical mapping showing conduction block after local hyperkalaemia. (C) Optical mapping showing preferential recovery of conduction through the previously inexcitable ischaemic zone along the reperfused artery. (D) Fluorescent beads showing local coronary vasculature. (E) Pre-hypoxia and (F) hypoxic block. (G) Hypoxia-reperfusion and (H) associated vascular bed. Acidosis does not gives rise to block of excitation, therefore no PVET nor re-entry. (I) Incidences of block, PVET and re-entries normalised to total performed experiments. SI n=39, hyperkalaemia n=10, hypoxia n=10 and acidosis n=12.

### Larger Ischaemic Size as a Predictor of PVET-based Arrhythmias

Stable conduction block and therefore PVET or re-entry during NF ischaemia-reperfusion was less likely in smaller hearts than larger hearts (Fig. 5B). The same was true in LoO_2_-reperfusion conditions (although in smaller hearts no metabolic inhibitor was used; Suppl. Fig. 3A). During HiK^+^-reperfusion, we increased the K^+^ concentration from 15 mM to 25 mM in order to achieve block in the smaller rabbits (Suppl. Fig. 3A). As a proof of principle, under this higher than standard K^+^ condition, it was possible to observe PVET and PVET-based re-entries in smaller hearts. As one would expect, the heart volume (Fig. 5C; 10.43 ± 0.321 mL, *N*=46 *vs* 5.844 ± 0.2571 mL, *N*=16; p<0.0001), weight (Fig. 5D; 9.58 ± 0.2311 g, *N*=46 *vs* 5.976 ± 0.236 g, *N*=14; p<0.0001), total rabbit body weight (Fig. 5E; 3155 ± 46.98 g, *N*=64 *vs* 1473 ± 41.86 g, *N*=20; p<0.0001) and heart to body weight ratios (Fig. 5F; 0.003096 ± 6.239e-005, *N*=47 *vs* 0.003816 ± 0.000223, *N*=11; p<0.0001) significantly differed between 4 and 2 month old rabbits, respectively.

### Reperfusion Paradigms

Finally, different reperfusion approaches to avoid PVET-induced re-entry were experimentally tested. As hyperkalaemia seemed to be the main driver of PVET-induced re-entry, followed by hypoxia, we tested a strategy of stepwise removing these *i.e.* ischaemia → hypoxia + acidosis → acidosis → physiological. Experimental testing of this showed no reduction in PVET or re-entry (Fig. 6F, *N*=5; block 100%, PVET 100% and re-entry 60%). We then focussed on HiK^+^ by testing a stepwise reduction in [K^+^]_o_ (15 mM, 13 mM, 10 mM, 6 mM; Fig. 6F, *N*=6; block 100%, PVET 83.3% and re-entry 50%), which was similar to direct one-step reperfusion (except for re-entry at 33.3%). Gradually reducing [K^+^]_o_ from 15 mM to 6 mM over 10 min of reperfusion resulted in no improvement either (Fig. 6F, *N*=3 block 100%, PVET 100% and re-entry 66.7%) compared to re-entry one-step at 33.3%. Also, a gradual transition from SI → HT (by gradual mixing of solutions) showed fewer PVET events (60%) but did not reduce re-entry (60%) (Fig. 6F, *N*=5). The only approach to reduce arrhythmias was by performing two-step reperfusion (we coin as ‘step-wise haemodynamic recovery of the ischaemic myocardium to lessen deleterious effects on cardiac electrophysiology’; SHIELD-EP). In the first step, the local catheter was advanced to perfuse only the distal area of the ischaemic region. After 10 min of distal reperfusion, to ensure the distal tissue had recovered excitability, the catheter was retracted to the occlusion site and reperfusion of the proximal site commenced. This approach resulted in shortened PVET paths that could not sustain arrhythmias (Fig. 6D, F; block 100%, PVET 87.5% but no re-entry).

**Figure 5:**
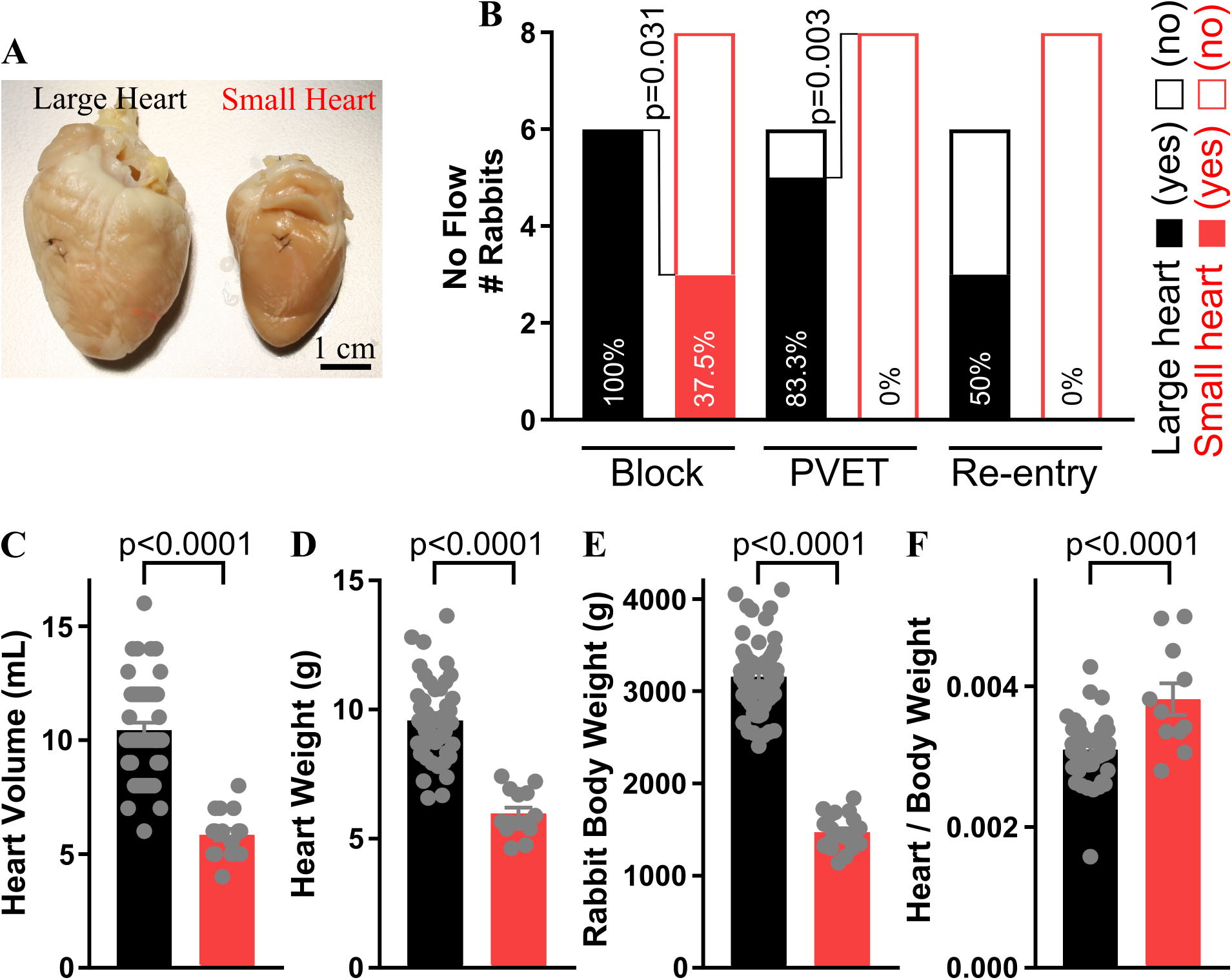
Incidences of PVET-based arrhythmias in larger *vs.* smaller hearts. (A) Comparing incidences of block, PVET and re-entry in no flow ischaemia-reperfusion in larger (black) *vs.* small (red) hearts (small hearts: NF n=8 *vs.* large hearts: NF n=6). (C-F) Characterization of small *vs.* large hearts volume, weight, animal weight and heart to body weight ratio.

**Figure 6:**
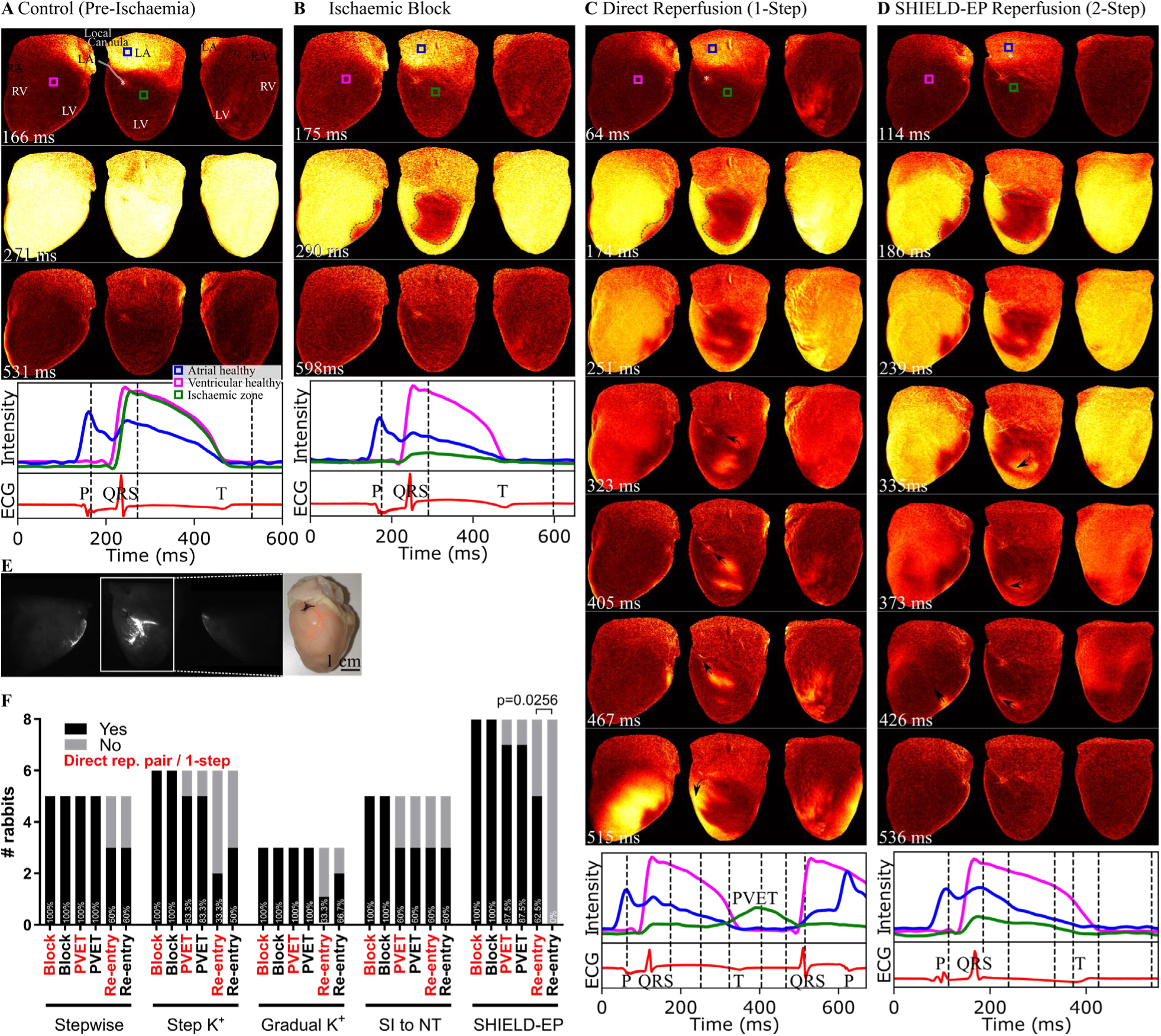
Two-step SHIELD-EP reperfusion approach shields from PVET arrhythmias. (A) Pre-ischaemic and (B) ischaemic block baselines. (C) Pre-intervention direct reperfusion showing PVET and associated PVET-based arrhythmias. (D) Using a two-step reperfusion approach, the proximal area is initially shielded from PVET-based re-entries. (E) Local beads in SHIELD-EP experiment. (F) Incidences of block, PVET and re-entries normalised to total performed experiments. All step-wise or gradual reperfusion strategy does not prevent PVET-based arrhythmias from the occlusion site. Stepwise reperfusion n=5, step K^+^ n=6, gradual K^+^ n=4, gradual SI-HT n=5 and SHIELD-EP n=8.

**Figure 7:**
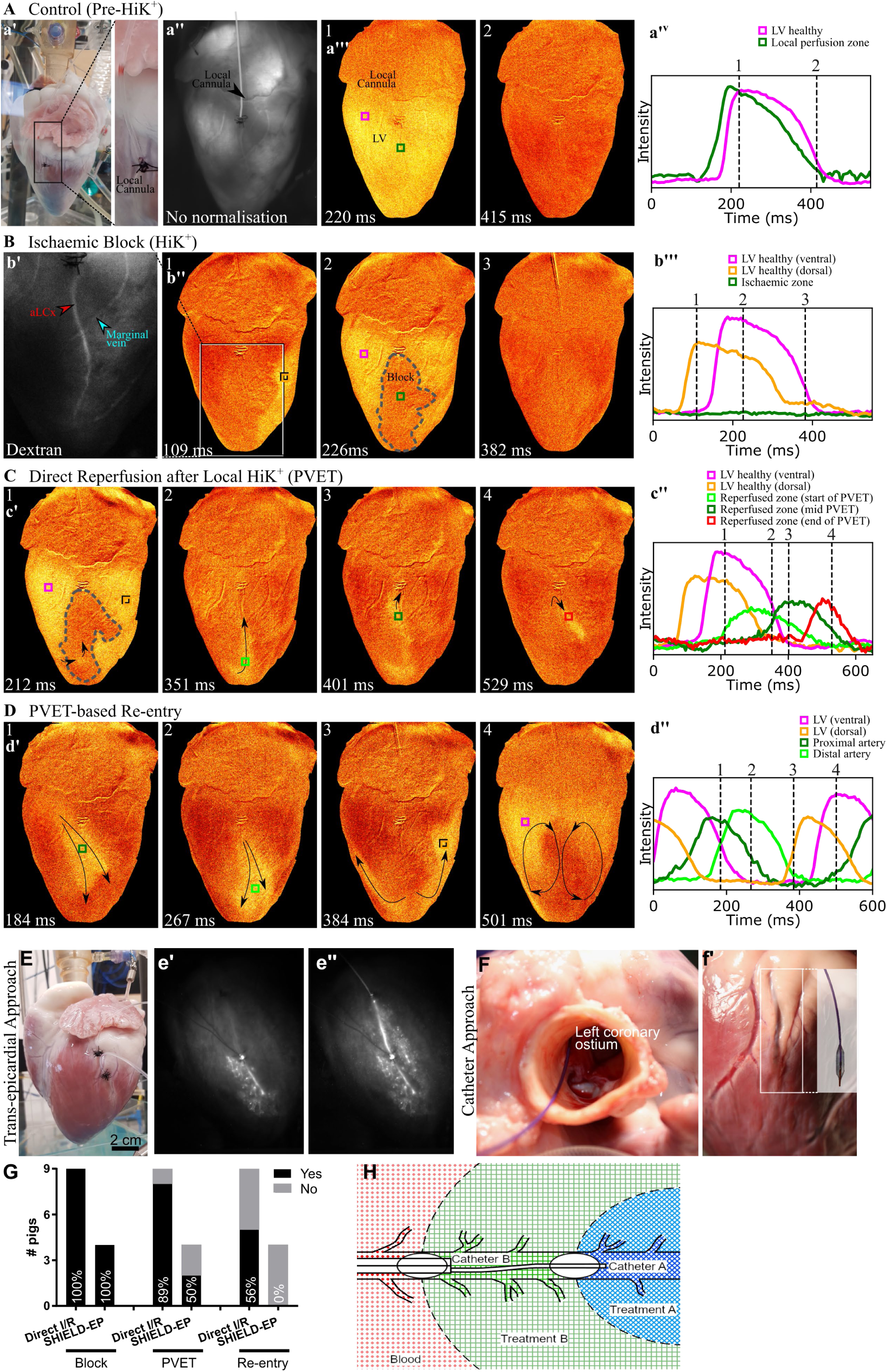
Translation evidence for PVET and protective SHIELD-EP. (A-D) Direct I/R n=9 and (G) SHIELD-EP n=4. Langendorff-perfusion with two transepicardial local coronary infusion sites (a’, E); inset shows distal first perfusion (e’) followed by proximal local perfusion (e’’) of fluorescent beads. Catheter prototypes must be <2mm (or 6F). Proof of principle balloon catheter insertion (F) through the coronary ostium in an *ex vivo* Langendorf perfusion-ready pig heart. Tailor-made aortic cannula to combine Langendorff-perfusion with catheter sheath access. Single balloon catheter test (f’) and design of possible irrigation-tip of patented dual-ballon / dual-perfusion-zone catheter (H) schematic of SHIELD-EP. The proximal balloon B is inflated upstream of the coronary occlusion, and a second, thrombuscrossing ‘distal balloon A catheter’ is pushed through and inflated downstream of the occlusion. Distal tissue re-perfusion with oxygenated physiological solution starts immediately, while the thrombus is removed. After removal, proximal perfusion starts, using oxygenated cardioplegic solution, to support re-oxygenation but not yet re-excitation. The distal catheter has electrodes to monitor electrical activity. Once distal excitation has normalised, the distal balloon is deflated and removed, while proximal perfusion with oxygenated physiological solution is started. Following full reperfusion, the proximal balloon is deflated and catheters are withdrawn.

### Translation

Final experiments involved moving from rabbits to a pig model to assess if PVET exists in a human-sized heart and if SHIELD-EP can prevent PVET-based re-entry. We observed PVET upon reperfusion of the ischaemic tissue with HT solution (Fig. 7C). This allowed excitation to enter and traverse the previously non-excitable ischaemic centre, giving rise to re-entry (Fig. 7D,G; direct I/R (*N*=9): block 100%, PVET 89%, re-entry 56%). PVET-based re-entry could be entirely circumvented via the SHIELD-ED approach (Fig. 7E,G,H (*N*=4): block 100%, PVET 50%, re-entry 0%).

## Discussion

Here, we identify PVET as a novel reperfusion arrhythmia mechanism following acute ischaemia. Not only do we describe a new mechanism, but we also offer a means of circumvent this arrhythmogenic substrate in pre-clinical and translational models.

### Experimental Evidence for Perivascular Excitation Tunnelling

The panoramic OM was essential for identifying PVET and earliest ‘break-through’ excitation by allowing visualisation of the entire cardiac surface. For the first time, we have demonstrated the early restoration of myocardial excitability along the trunk of the reperfused coronary artery. This forms a perivascular tunnel with slow conduction that traverses the ischaemic core, providing a path for re-entrant arrhythmias. Re-entry may progress in a baso-apical (descending) or apico-basal (ascending) direction, likely due to differences in Purkinje activation pattern, vessel morphology, action potential duration, and speed of conduction in physiological and ischaemic tissue.

The return to normal physiological conditions after a period of local ischaemia generates EP heterogeneities due to differential reperfusion of the affected tissue. OM recordings showed a preferential return of conduction, primarily along the main artery trunk, where the tissue was reperfused earlier. There was no difference in the return of conduction between no flow I/R and SI-reperfusion (Fig. 2), with both procedures leading to similar incidences of PVET and re-entry. This suggests that PVET may be primarily driven by the effects produced by the heterogeneous return to normal physiological [K^+^]_o_, pH, and pO_2_, instead of changes in gap junction coupling or cell damage.

### Teasing out Key Drivers of PVET

PVET appears to be largely a physical phenomenon driven by the architecture of the coronary vasculature. The main condition necessary for PVET to occur is that a sufficiently large conduction block is present through which the perivascular excitation can travel. Individually dissecting ischaemia by the three main factors in the acute phase (*i.e.*, hyperkalaemia, hypoxia, and acidosis) showed that heterogeneities in pH of the tissue do not lead to conduction block, therefore no PVET or re-entry. On the contrary, reperfusion from locally elevated K^+^ and the decrease of metabolic activity resulted in PVET and re-entrant arrhythmias. Conduction block is produced in hyperkalaemia and hypoxia by an elevation of the transmembrane voltage (V_m_), that inactivates Na^+^ channels. The elevation of V_m_ is likely caused in both cases by an imbalance of the physiologic intra- and extracellular [K^+^]. The Nernst potential of K^+^ is directly proportional to log([K^+^]_o_/[K^+^]_i_), therefore in HiK^+^, the increase in [K^+^]_o_ drives V_m_ to a more positive value. In the case of hypoxia, we hypothesise that the impairment of the Na^+^-K^+^ pump and the activation of the K-ATP channel, produces a depletion of intracellular K^+^, which has the same effect on the Nernst potential of K^+^ as hyperkalaemia. In contrast, acidosis does not have a noticeable effect on V_m_ nor any other mechanism to achieve conduction block that could produce the substrate for PVET in our simulations.

When considering PVET-based arrhythmia incidence following conduction block, hyperkalaemia-reperfusion results in the highest likelihood of PVET and re-entries. This is in line with previous studies showing that K^+^ heterogeneities alone are sufficient to provide the substrate and the trigger for re-entrant ventricular fibrillation during reperfusion.^24,25^ It furthermore, is in line with the relative effects of changes of intracellular and extracellular potassium on the Nernst potential, where mM level increases in extracellular potassium can cause a dramatic rise in the Nernst potential, whereas order of magnitude greater changes are needed intracellularly to achieve a similar effect.

### Anatomical Substrate: Coronary Artery Morphology is a Determinant of PVET

It is known that the adventitia surrounding major coronary arteries insulates the extracellular space, and exchange between blood and extracellular fluid is at the capillary level. Our data on coronary flow distribution suggest that the myocardium closest to the main coronary artery trunk is ‘privileged’ in terms of speed of access to coronary perfusate (Fig. 3). This provides a new look at the underlying anatomical substrate of the surprisingly fast recovery of excitability in perivascular myocardium.

Variability in the coronary vasculature leads to different presentations of PVET and the kinetics of both conduction block and full recovery. PVET presents itself in different morphologies, depending on the out-branching degree of the cannulated aLCx. Moreover, the size of the affected region as determined by the location of the ‘block’ and the vascular network is a decisive factor in whether PVET may cause re-entry or not, since this type of arrhythmia is highly dependent on the length of the re-entrant path (and, therefore, dependent on the size of the ischaemic area). Thus, a larger affected area will be more prone to re-entry (Fig. 5). The rabbit model has little collaterisation between arteries, similar to both pigs and humans.^26^ Given the dependence of PVET on coronary architecture, it may be that assessment of the left coronary artery branching pattern and/or flow bed may allow prediction of arrhythmia susceptibility after coronary occlusion / reperfusion (as shown for chronic heart failure models).^27^

### How to Prevent PVET-based Arrhythmias?

A number of reperfusion strategies for preventing PVET and PVET-based arrhythmias were tested. None of those which reperfused directly from the occlusion site prevented PVET and PVET-based re-entry; if anything, the rates were increased. This is likely due to these reperfusion strategies extending the duration where there were significant EP gradients. When perfusing with a potassium concentration slightly below that which caused complete block, we observed areas where block had already occurred and other areas that remained excitable (data not shown), probably due to a combination of potassium gradients between globally and locally perfused tissue and due to different source-sink relations across the tissue area. These ‘natural’ heterogeneities in EP mean that PVET may be an unavoidable phenomenon and so strategies should instead focus on preventing PVET-induced arrhythmias.

Controlling the size of the reperfusion area prevented arrhythmias. This was achieved through our SHIELD-EP approach. By advancing the local cannula deep into the tissue to rescue the distal part first followed by proximal reperfusion we were able to prevent PVET-based arrhythmias. PVET-based arrhythmias were prevented in the first step by blocking their re-entry path (the inexcitable proximal tissue acting as a shield) and in the second step by reducing the PVET path length to the extent that re-entries could not occur.

### Limitations

Experiments were performed on hearts from relatively young and healthy animals without risk factors, comorbidities and co-medications (as would be characteristic of patients suffering from myocardial ischaemia).^28^ While it would be desirable to perform experiments in aged rabbits, this is not feasible, as New Zealand White rabbits live for up to 10 years.^29^ Our experiments may therefore underestimate the relevance of PVET. This will form a target of subsequent studies in aged/diseased models. We do not expect sex-differences at these juvenile ages.

Another limitation was the approach of making a small incision on the epicardial surface to insert the local cannula, rather than non-invasively guiding the custom plastic catheter through the left coronary artery ostia (*e.g.* Fig. 7F). Such an elegant approach would be possible in a larger animal model, evading potential leakiness from cutting into the aLCx or unintended tissue damage to surrounding vessels, nerves or other structures. However, while single balloon catheters exist, they do not facilitate local perfusion as needed for a non-invasive SHIELD-EP approach.

Optimal spatio-temporal resolution of cardiac OM is usually achieved in the presence of pharmacological excitation-contraction uncouplers such as blebbistatin. However, AP duration in mechanically-arrested Langendorff-perfused rabbit hearts is 25% longer than in contracting preparations, and ventricular fibrillation inducibility is significantly reduced in the presence of blebbistatin.^30,31^ This means that our experiments may be underestimating the likelihood of PVET-driven arrhythmogenesis. Additional experiments should compare PVET occurrence in uncoupled *vs*. beating hearts. In future, post-processing of OM recordings with motion tracking algorithms should be used to reduce motion-related artefacts allowing experiments to be performed in freely-beating hearts.^31^ Crystalloid perfusates tend to cause tissue swelling (due to low oncotic pressure), and media free of blood cells are not ideal for preparations with high oxygen demand.^32^ However, as our experiments were performed on mechanically uncoupled hearts, this was not problematic, nevertheless in future working heart experiments, blood supplementation may be necessary.

### Clinical Relevance & Outlook

In the clinical setting, catheter-based reopening of blocked coronary arteries (percutaneous coronary intervention or PCI) is commonly used in patients with myocardial ischaemia^33^ to recover blood supply, reduce infarct size and preserve heart function.^34^ It is plausible that PVET occurs during reperfusion also in patients (though currently difficult to delineate with clinically available tools). It may be possible to reduce IRA risk for patients undergoing primary percutaneous coronary intervention^34,35^ using an irrigated catheter to flush the previously ischaemic zone upon reperfusion with specifically designed solutions, prior to restoring normal blood flow. Developing our SHIELD-EP 2-step reperfusion strategy that prevents the occurrence (clinically and/or duration) of PVET-induced arrhythmias would be of significant translational relevance (Fig. 7H).

All in all, PVET offers a novel conceptual explanation for reperfusion arrhythmias with now known drivers and a means of preventing PVET-induced re-entry with the hope to eventually reduce IRA burden upon cardiac revascularisation in patients. Our ongoing work will apply SHIELD-EP in clinical trials once we develop a coronary balloon catheter to support a minimally-invasive route to testing SHIELD-EP *in vivo*.

So what? In Germany, >300,000 patients undergo PCI and in the USA >900,000 per annum. IRA following PCI is seen in ∼5% (some reports quote up to 23%) of cases^36–39^. Defibrillation, systemic thrombolytics, and conventional PCI are good, but better therapies are needed. The clinical opportunity for the SHIELD-EP approach would thus range from 15,000 to 69,000 cases p.a. in Germany alone. Our approach may take the ∼86% successful PCI to 100% with SHIELD-EP.

## Acknowledgments

We thank Prof Constantin von zur Mühlen and Mr Thomas Kok for the translational work support and Dr Gunnar Seemann for early project guidance. Also, we thank the Freiburg Cardiovascular Biobank (CVBB) via Ms Stefanie Perez Feliz, Dr Simone Nübling, Mr Jonas Heer, Mr Manuel Koch, Ms Pia Iaconianni, Mrs Cinthia Buchmann and Ms Kristina Kollmar for excellent technical assistance. Much thanks is given to the Chair of Livestock Biotechnology, Prof. Angelika Schnieke and Dr Konrad Fischer, Technical Univerisity of Munich (TUM), WZW Weihenstephan, Liesel-Beckmann Str, 1, D-85354 Freising for the hybrid pig model support. Locally, a special thanks to Dr Heidi Ramona Cristina Schmitz and Mr Johannes Dinkelaker from CEMT-Freiburg Experimental surgery for large animal handling support. We thanks Prof Christof Benk for the ECMO donation and Mr Thomas Beha for the Hugo Sachs technical support.

## Abbreviations

aLCx: Anterior branch of the left circumflex coronary artery AP Action potential(s)
ATP: Adenosine triphosphate
EP: Electrophysiolog(y/ical)
HiH^+^: Acidosis condition
HiK^+^: Elevated extracelular potassium condition
HT: Oxygenated HEPES-buffered Tyrode’s solution
IRA: Ischaemia-reperfusion arrhythmia(s)
[K^+^]_o_: Extracellular potassium concentration
LoO_2_: Reduced oxygen partial pressure and metabolic inhibition condition NF No flow
OM: Optical mapping
PCI: Percutaneous coronary intervention
PVET: Perivascular excitation tunnelling
SHIELD-EP: Step-wise haemodynamic recovery of the ischaemic myocardium to lessen deleterious effects on cardiac electrophysiology
SI: Simulated ischaemia solution
V_m_: Transmembrane voltage

