## Supplemental Figures for "Perivascular Excitation Tunnelling: a Novel and Preventable Cause of Cardiac Reperfusion Arrhythmias"

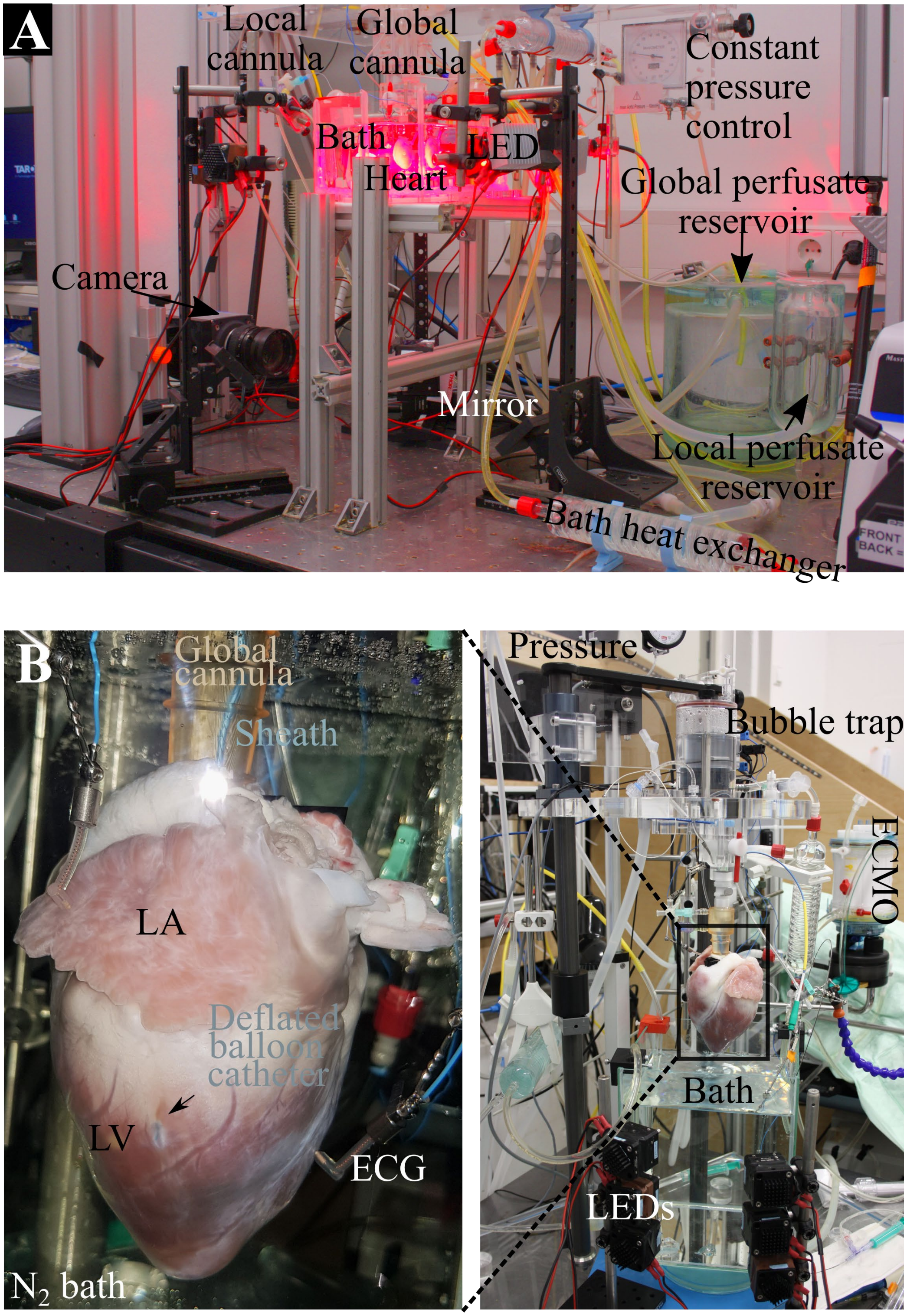


**Supplemental Figure 1: Photograph of the experimental setup** (components labelled) of panaromic rabbit hardware (A) and single view pig optical mapping (B).

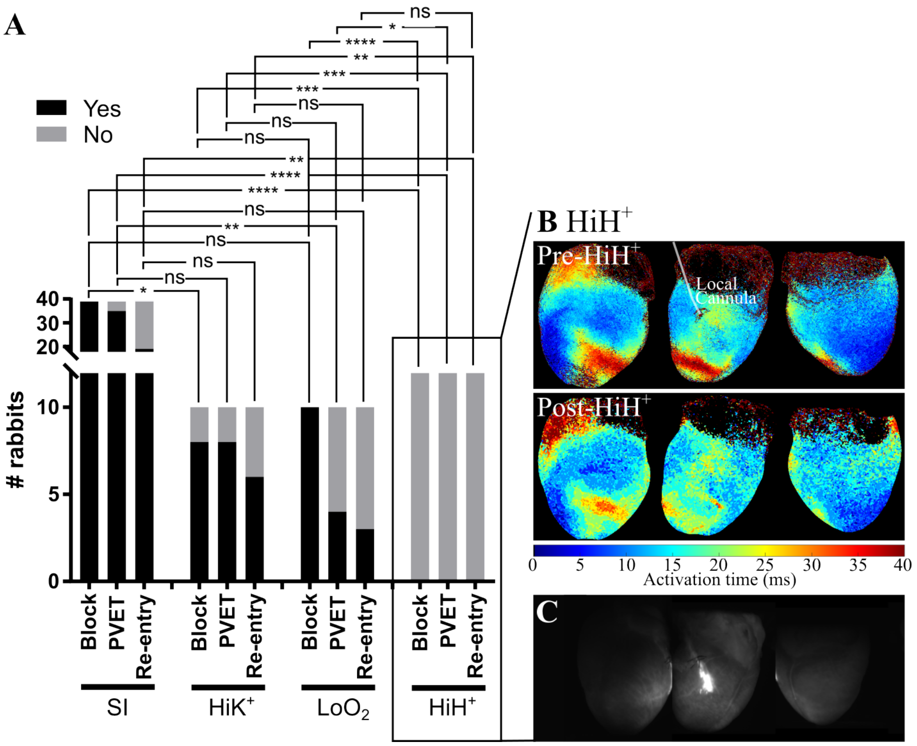


**Supplemental Figure 2: Acidosis-Reperfusion.** Acidosis does not gives rise to block of excitation, but a slight slowing of activation after 1 hr of local acidosis (B). Local beads in acidosis experiment (C). Incidences of block, PVET and re-entries normalised to total performed experiments. SI n=39, hyperkalaemia n=10, hypoxia n=10 and acidosis n=12.

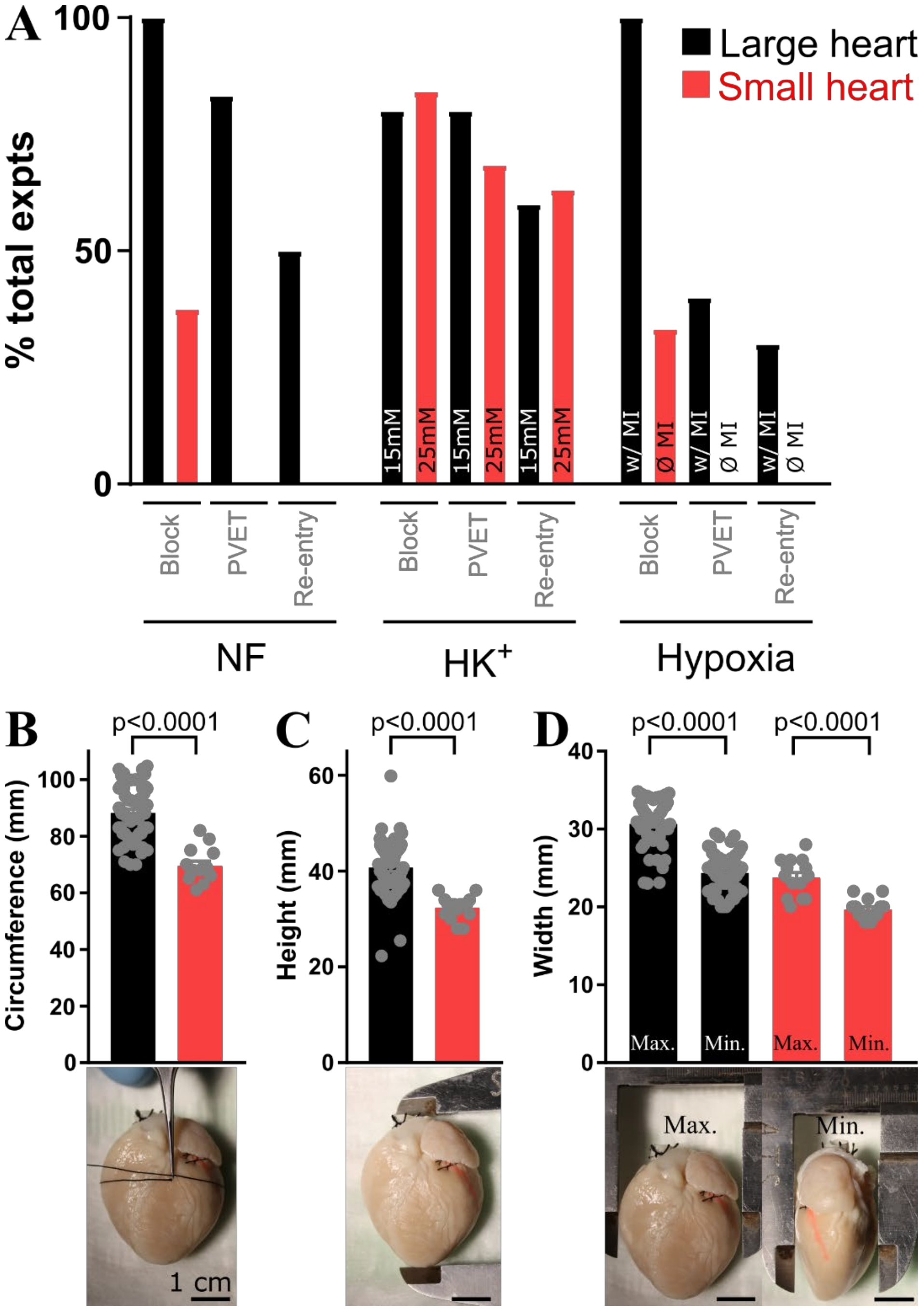


**Supplemental Figure 3: Incidences of PVET-based arrhythmias in larger *vs.* smaller hearts.** (A) Comparing incidences of block, PVET and re-entry in no flow ischaemia-reperfusion in larger (black) vs small (red) hearts (small hearts: NF n=8, HK [25mM] n=19, hypoxia n=3 *vs.* large hearts: NF n=6, HK [15mM] n=10, hypoxia+MI n=10). Harder to achieve block in small heart NF-reperfusion and so no PVET or re-entry *vs.* in larger hearts. In small heart during hypoxia-reperfusion (no metabolic inhibitors used), fewer hearts reached ischaemic block therefore no PVET or re-entries were observed. In smaller hearts where a higher concentration [25 mM instead of normal 15 mM] of $K^+$ was perfused, PVET and PVET-based arrhythmias can be observed. (B-D) Characterization of small *vs.* large hearts circumference, height, and width.
